# Insights into the successful breeding of Hawksbill sea turtles (*Eretmochelys imbricata*) from a long-term captive breeding program

**DOI:** 10.1101/2020.06.26.174219

**Authors:** Ruth Maggeni, William E Feeney

**Affiliations:** Environmental Futures Research Institute, Griffith University, Nathan QLD 4111, Australia; Department of Behavioural Ecology and Evolutionary Genetics, Max Planck Institute for Ornithology, Seewiesen, Germany; Department of Zoology, University of Cambridge, Cambridge CB23EJ, United Kingdom

**Keywords:** Captive breeding, Conservation, *Eretmochelys imbricate*, Hawksbill sea turtle, Sea turtle

## Abstract

Sea turtle populations are declining and evidence-based methods for supporting their populations are required. Captive breeding and release programs can be effective, offering the opportunity to supplement nature populations; however, sea turtles require specific conditions to successfully breed. Here, we present insights gained from a 16 year Hawksbill sea turtle (*Eretmochelys imbricata*) captive breeding program that was conducted at the Underwater Observatory Marine Park, Eilat, Israel, between 1982-1997. As the first program of its kind for the Hawksbill sea turtle, insights were gained largely through trial-and-error and word of mouth advice. The key insight gained during this program was the critical importance of pre-breeding separation of the sexes; turtles did not breed prior to pre-breeding separation being implemented, but it became predictably regular once it was. Over the course of the program, 161 two-three years old hatchlings were introduced to the Red Sea, which was enabled largely as a result of pre-breeding separation being implemented.

## 1. Introduction

Sea turtles (Chelonioidea) are long-lived migratory species that inhabit almost all of the world’s oceans (Wallace et al. 2011). Seven species exist, with three being listed as Critically Endangered (Hawksbill, *Eretmochelys imbricata*; Kemp’s ridley, *Lepidochelys kempii*; and leatherback, *Dermochelys coriacea*), one listed as Endangered (green, *Chelonia mydas*), two listed as vulnerable (loggerhead, *Caretta caretta*; and olive ridley, *L. olivacea*) and one as Data Deficient (flatback, *Natator depressus*) according to the Red List (IUCN 2020). Major threats to sea turtle populations include being hunted for their eggs, meat, or shells; being caught as bycatch; and habitat destruction (Wallace et al. 2013; Lewison et al. 2014; D’Cruze et al. 2015; Mazaris et al. 2017). All species are listed as having decreasing population trends, with exception of the flatback and Kemp’s ridley, for which their population status’ are unknown (IUCN 2020). Worryingly, the general lack of data and unevenness of past research efforts is a key concern for the management of sea turtle populations globally (Rees et al. 2016; Rees et al. 2018).

The biases of past research mean that the assessment of less studied populations or species risk falling victim to “shifting baseline syndrome” (Sheppard 1995). For example, while the green and loggerheads sea turtles have received the overwhelming majority of past research attention (41% and 34% of publications, respectively), those that are under significant threat (i.e. Critically Endangered), such as the Hawksbill, Kemp’s ridley and leatherback turtles together were the focus of only 5% of papers (Rees et al. 2016). Concerning the Hawksbill sea turtle, the only available data regarding population changes dates back two generations (i.e. a few decades), which makes it difficult to assess how human activities have damaged their populations (Meylan 1999). Thus, given that past research efforts have been patchily distributed and that management of long-distance migrants, such as sea turtles, presents inherent challenges (Campbell 2007; Jeffers and Godley 2016), the preservation of sea turtle populations and species requires the use of additional conservation alternatives.

Several alternative conservation measures are currently being used for the Hawksbill sea turtle, which vary in their aims and effectiveness. For example, longstanding public information campaigns in countries including Samoa and Brazil have highlight the ecological importance of this species as well as the risks posed to it by human activities (Witzell 1974; Marcovaldi and Dei Marcovaldi 1999). Population monitoring, through the use of satellites is also being used throughout regions including central Africa and the eastern Mediterranean to elucidate migration patterns (Broderick et al. 2007; Maxwell et al. 2011). Like other sea turtles, hatching success and juvenile mortality represent potential population bottlenecks, and long-term “head starting” programs have been running in the Cayman Islands and Florida (Bell et al. 2005). Head starting is where hatchlings are transferred into hatcheries, and later kept in a protected environment for 9-12 month, until they are released into the sea, providing them with higher chances of survival at an older age (Bell et al. 2005). In 1971 the fisheries division of western Samoa collected 500 eggs in each nesting season and transferred them into hatcheries where they were kept to protect hatchlings (Witzell 1974). Previous attempts of captive breeding showed females produce two to five times more eggs in captivity than in the wild, a factor which could be highly significant in the rehabilitation of a population (Wood et al. 1980). The potential effectiveness of such captive breeding and re-introduction programs is highlighted by the Cayman Turtle Centre LTD, which has been operational in the Cayman Islands since 1968, and 90% of the wild population of green turtles can be traced back to the population of the turtles in the farm (Barbanti et al. 2019).

In this study, we report the insights gained from a 16 year Hawksbill sea turtle captive breeding project that was operated by Coral World Eilat, Israel, between 1982 and 1997.

## 2. Materials and methods

### 2.1 Study site and rebreeding program

The breeding program took place over a 16 year period at the Underwater Observatory Marine Park Eilat, Israel, between 1982-1997. Four female Hawksbill sea turtles (*hereafter* “turtles”) were collected from the south Red Sea (28.2439, 34.4674) age 8-10 years old and 8 months later one 10 year old male was collected from Tiran (28.0189, 34.4753). The turtles were housed in a 40×20m 3m deep pool with a 15×10m 1m deep sand island that extended from one side of the pool to its centre. Thus, the turtles had unlimited access to the sand island from three different directions. The sand was collected from the nearby beach at (29.5033, 34.9182), as required. While held in captivity, the turtles were exposed to natural light conditions and were fed a diet of lettuce and fish twice a day. Animal care and use guidelines of the Coral World – Marine Parks Worldwide were followed by all facility operators. The sand island was monitored 24 hours per day and any mating behaviours were noted by the facility operators (described in detail below). Once eggs were laid, the site was fenced to prevent other turtles from disturbing the nest. The humidity and temperature of the nest was also measured twice per day (day and night), using sensors and thermometers.

### 2.2 Animal behaviour and care

Mating behaviours were recorded as indicators of sexual arousal in males, possible fertilization in females, and highlighted that the conditions were satisfactory as a suitable breeding environment. These behaviours included the males biting the female on her neck and rear flippers as well as nuzzling her head. Copulation occurred following these behaviours if the female made no attempt to distance herself from the male. Avoidance of the male by the female was classified as a negative interaction. After observing no evidence of mating behaviours during years 2-5, it was suspected that the lack of interest by males was due to them being housed with females throughout the year, with no time apart. It was therefore decided to try separating the males and females at the end of each mating season until the beginning of the following mating season. After seeing successful mating events in years 6 and 7, we decided to introduce control groups in each subsequent year to investigate the role pre-mating separation played in encouraging successful mating. Therefore, in year 8 through 12, each year included a control group with similar number of males and females, where no separation took place throughout the mating season. All other conditions of the control group were identical to the separated group.

Once a female laid eggs, the eggs were then immediately fenced off (year 1, 6-7), to ensure their protection until they hatched. In years 8-12 eggs were transferred to an incubator. This decision was made as incubators provide more controlled environmental conditions (i.e. humidity and temperature) to help ensure that healthy hatchlings were produced. Initially, two temperature groups were created in the incubators. First, 27°-29° to produce male turtles, and 28°-32° to produce female turtles. Ventilators and sprinklers managed the humidity in the incubators and different humidity levels were tested regularly.

Once hatching began, hatchlings were collected and examined to ensure they were healthy. They were also measured and weighed to allow the curators to document their development. For the first two years of their life, newly hatched turtles were kept in 1.5*1m plastic containers, which contained aired sea water. During this time the hatchlings were handfed fish using tweezers three times per day. Hatchlings were also weighed and measured once per week to monitor growth rate. Between the age of 2-8, up to 10 juvenile turtles were kept in 2*1.5m deep pools. The pools contained natural characteristics such as rocks, seaweed, and some reef fish. Similar to adult turtles, juveniles were fed twice a day with chopped fish and whole lettuces. Juvenile sea turtles were weighed and measured once a month.

### 2.3 Introduction to the Red Sea – selection and handling

After being raised for two-three years in the aquarium, hatchlings went through different tests in order to ensure their health and their ability to survive in natural conditions prior to their introduction to the sea. The tests included measurements (e.g. length, width, weight and temperature, which was measured by inserting a thermometer 7-10 cm inside the cloaca) The hatchlings were also introduced to live pray such as crustaceans and jellyfish, and hatchlings which demonstrated the ability to catch live pray were considered fit for survival in natural conditions. Some healthy turtles were kept to help support future breeding years. Once the decision to release the turtles into the sea was made, the turtles were placed in smaller containers and carried to a beach (29.5033, 34.9182), which is closed to the public and therefore provided a safe environment along with a reach marine environment. Then, along with a curator snorkelling by, each turtle was escorted at arm’s length distance until both the curator and the turtle reached deep water (approximately 6m) at which point the curator allowed the turtle to advance on its own. After five years the area of introduction was expanded to an area up to 50km away from the original release site using a boat.

### 2.4 Statistical analysis

Analyses were conducted in R statistical software v1.1.4 (R Core Team 2020). Data used in this manuscript are provided as Supplementary Information.

## 3. Results

At the start of the project (year 1, 1982), limited mating behaviour was observed, which produced 88 eggs (22 ± 7.39 eggs per female mating opportunity, range = 0-32, *N* = 4 females). Of the three turtles that produced eggs, one female produced five hatchlings from 32 eggs and the other two produced no hatchlings. Inspection of the eggs indicated that these eggs were under-developed. During years 2-5 (1983-1986) the males showed no interest in the females and no mating behaviours or mating events were recorded (0 eggs produced by 29 female mating opportunities [i.e. female identity was not recorded across years] comprising 6-8 females per year). Successful mating was observed when pre-breeding separation was implemented in years 6-7 (1987-1988) (41.08 ± 3.54 eggs per female mating opportunity, range = 30-65, *N* = 13 female mating opportunities). A control treatment (i.e. no pre-breeding separation) was implemented from year 8-12 (1990-1992 and 1995-1996), which was associated with a drastic increase in mating behaviours and egg production by females (43.87 ± 3.29 eggs per female mating opportunity in the pre-mating separation group, range = 0-70, *N* = 45 female mating opportunities, compared to 0 eggs per female mating opportunity in the control group, *N* = 34 female mating opportunities). We also found no difference in the hatching success between eggs incubated in an incubator (years 8-12, 24.82 ± 2.16 hatchlings per female mating opportunity [56.59% hatch rate], range = 0-45, *N* = 45 female mating opportunities) compared to when they were left to incubate in the sand (years 1, 6-7, 21.92 ± 2.36 hatchlings per female mating opportunity [53.37% hatch rate], range = 16-41, *N* = 13 female mating opportunities) (t-test: *t* = 0.91, *df* = 33.9, *P* = 0.37).

## 4. Discussion

Twelve years of Hawksbill captive breeding was conducted at the Underwater Observatory Marine Park Eilat between 1982-1997. As the first Hawksbill captive breeding program of its kind (at least as far as the Marine Park staff understood), the eventual successful captive breeding of turtles was a product of trial and error that was informed by word-of-mouth knowledge. Several key insights were made, most notably the important role that pre-breeding separation of male and female turtles had on males exhibiting interest in females and successful mating events. Additional insights, such as potential usefulness of local knowledge and the relative hatching success of eggs incubated in sand versus an incubator were also gained.

Our data highlight the importance of pre-breeding separation for successful breeding of Hawksbill sea turtles in captivity. Sea turtles are naturally a solitary species outside of the nesting season (Lund 1985; Stringell et al. 2013), and thus separating them prior to breeding is intuitive as it resembles natural conditions. As such, captive breeding programs have reported/recommended the use of pre-breeding separations (e.g. Green sea turtles, Wood et al. 1980); however, why this practice is used and empirical data regarding its impact remains unreported. Likewise, while pre-breeding separation is common across a wide array of species (e.g. southern corroboree frog, *Pseudophryne corroboree* (McFadden et al. 2013), corn snakes, *Elaphe guttata guttata* (Griswold 2001), cobia fish, *Rachycentron canadum* (Gopakumar et al. 2011), mallards, *Anas platyrhynchos* (Stunden et al. 1999) and primates (Wallace et al. 2016), surprisingly few studies have experimentally investigated the effect this practice has (for a notable exception, see McFadden et al 2013). Our data indicate that pre-breeding separation is vital for successful mating, with no mating events recorded in the four years separation was not implemented, compared to an average of ∼44 eggs per female following its implementation.

The information gained through this study also highlights the value of local word-of-mouth information for captive breeding programs. To our knowledge, this project comprised the first effort to captively breed Hawksbill sea turtles for the purposes of reintroduction to their natural environment. Given that there was little literature available to us at the time, we sought help from potentially informed individuals to help overcome the lack of breeding observed in years two-five of the program. In 1986, we reached out to a Hawaiian sea farmer in Maui, who kept a small group of Hawksbill sea turtles in the bay he lived in. The turtles were kept in a sheltered lagoon surrounded by nets, for the purpose of rehabilitation in a safe environment. The sea farmer suggested that we try to separate the males and females prior to the start of the mating season. He explained that spending an entire year in the same pool, the male turtles lose interest in the females and by the time mating season begins, the males show no interest in the females. This proved extremely successful and was an insight that underpinned the subsequent success of this program in the following years.

We found no evidence that incubators increased hatching success during this program. Once pre-breeding separation was implemented and females were consistently producing eggs, we consistently recorded an approximately 55% hatch rate, regardless of whether eggs were incubated within sand or an incubator. This contrasts with other, more recent, studies that have demonstrated hatch success rates of up to 90% when an incubator is used (Khalwatu 2020), which may be a result of more insights gained since our data was collected in the 1980s and 1990s. Incubators no doubt can be extremely useful when used to protect incubating eggs from predators (Wyneken et al. 1988), and to influence the sexes and condition of the resulting offspring (Hewavisenthi and Parmenter 2002; Reneker and Kamel 2016). In the case of this breeding project incubators were incorporated mainly to compensate for the lack of space on the sand island within the enclosure. Conditions on the sand island were relatively consistent throughout the duration of the incubation period and because the nests were fenced off, they were protected from any outside disturbance that could occur in natural conditions, which may help explain the similar hatch success of eggs in the sand versus the incubator.

Over the 16 years that this captive breeding program operated, successful breeding was recorded in seven years and 161 Hawksbill sea turtles were introduced to the sea following two years in the aquarium. As the first project of its kind for this species, insights were hard won, primarily through trial and error, as well as through word-of-mouth information gained from knowledgeable individuals. The key insight was the important role that pre-breeding separation plays in facilitating captive breeding, which while commonly used in taxonomically diverse captive-breeding programs has rarely been experimentally validated. Given that similar practices have been reported in other sea turtles, this study reiterates its usefulness to increase the effectiveness of these programs, which can play an important role in the management of these charismatic but at risk species.

## 5. Acknowledgements

We thank Aharon Miroz, Head Curator of Coral World International for access to these data. We would also like to thank the curators of The Underwater Observatory Marine Park – Eilat, Israel for their hard work which made this project successful.

## 6. Permits

Collection and captive breeding was conducted with permission from The Israel Nature and Parks Authority (1997/494).

## Notes

### Competing Interest Statement

The authors have declared no competing interest.

